# Focus: The interface between data collection and data processing in cryo-EM

**DOI:** 10.1101/105452

**Authors:** Nikhil Biyani, Ricardo D. Righetto, Robert McLeod, Daniel Caujolle-Bert, Daniel Castano-Diez, Kenneth N. Goldie, Henning Stahlberg

## Abstract

We present a new software package called *Focus* that interfaces cryo-transmission electron microscopy (cryo-EM) data collection with computer image processing. *Focus* creates a user-friendly environment to import and manage data recorded by direct electron detectors and perform elemental image processing tasks in a high-throughput manner while new data is being acquired at the microscope. It provides the functionality required to remotely monitor the progress of data collection and data processing, which is essential now that automation in cryo-EM allows a steady flow of images of single particles, two-dimensional crystals, or electron tomography data to be recorded in overnight sessions. The rapid detection of any errors that may occur greatly increases the productivity of recording sessions at the electron microscope.

**Abbreviations**
cryo-EM: Cryo-electron microscopy
DED: Direct electron detector
2D: Two dimensional
3D: Three dimensional
CTF: Contrast transfer function
FFT: Fast Fourier transform
GPU: Graphical processing unit
CPU: Central processing units
MRC: MRC file format
SNR: Signal-to-noise ratio
GUI: Graphical user interface

## 1. INTRODUCTION

Cryo-electron microscopy (cryo-EM) and cryo-electron tomography are employed in structural biology to determine the structure of vitrified biological samples, such as isolated proteins and protein complexes, two-dimensional (2D) protein/lipid crystals, or larger biological objects such as bacteria and nanocrystals. Until the development of direct electron detector cameras (DED) for electron microscopy, the resolution achieved in most cases was in the 5 - 30Å range, with few exceptions (Gonen et. al., 2005). Unlike detectors that convert the electron signal to light via a scintillator, DEDs have radiation-hardened metal oxide semiconductor sensors that can be directly illuminated with the high-voltage primary electron beam. This architecture greatly reduces the point-spread function of the electron signal and results in superior detector quantum efficiency, enhanced readout speed and an excellent signal-to-noise ratio (SNR). The resolution required to count single electron events is achieved. Furthermore, the low read-out noise of DEDs makes it possible to record electron dose-fractionated multi-frame exposures, known as image stacks or movies. This means that movement of the sample during imaging can be corrected by software, as discussed further below. Averaging the aligned motion-corrected frames after they have been weighted by a dose-dependent B-factor to account for damage to the proteins by the electron beam, results in images that have superior resolution and a better SNR (Rawson et al., 2016). Together with improved image processing algorithms, such as maximum-likelihood (Lyumkis et al., 2013) and Bayesian (Scheres, 2012) methods, this has lead to a resolution revolution in cryo-EM (Kühlbrandt, 2014) with the result that atomic models of the imaged biological structures can quite often be built (Crowther, 2016; Kühlbrandt, 2014; Liu et al., 2016;Merk et al., 2016;Yu et al., 2016).

The Gatan K2 Summit DED camera was the first to offer three recording modes. In the first mode, the linear mode, the energy deposited in the DED sensor by passing electrons is integrated over the entire exposure time. In the second mode, termed electron counting, single electron events are individually detected and counted. Finally, in the super-resolution mode, the camera driver registers the impact location of individual electrons on the detector with sub-pixel accuracy during the electron counting step, resulting in final images of twice the pixel resolution than the hard chip of the DED detector. Image data are ideally recorded in super-resolution mode as multi-frame exposures of each region of interest. Use of the super-resolution algorithm means that the “4k” chip (3838 × 3710 pixels) of a K2 Summit DED records “8k” frames (7676 × 7420 pixels). Subsequently downsampling these frames to “4k” by cropping in Fourier space, results in 4k images with a SNR significantly superior to the SNR of 4k images directly recorded in the “4k” counting mode, due to the effect of antialiasing (see alsoRuskin et al., 2013).

When multi-frame exposures are recorded using a DED, the total dose given to a region of interest is fractionated, allowing the images in the stack to be aligned by postprocessing to eliminate specimen movement originating from physical drift of the specimen stage, electronic drift in the imaging system of the microscope, or physical movement of the sample under the electron irradiation. Several software packages are available to correct specimen motion, including Zorro (McLeod et al., 2016), MotionCor2 (Zheng et al., 2016), Unblur (Grant and Grigorieff, 2015), alignparts_lmbfgs (Rubinstein and Brubaker, 2015), MotionCorr (Li et al., 2013), and SerialEM (Mastronarde, 2005). Among them, Zorro, MotionCor2 and alignparts_lmbfgs perform both whole-frame and local drift-correction. In most cases, additional local drift-correction significantly improves the quality of the output image, but also increases the computational costs considerably. Optimally, the microscope operator should be able to view drift-corrected and averaged images directly on the camera computer during the microscopy session. Thus, the development of software tools that can achieve whole-frame ‘on the fly’ drift correction has gained interest (Li et al., 2015; Nobel and Stagg, 2015;Fernandez-Leiro and Scheres, 2016). Local-drift correction cannot (yet) be computed during data collection on a conventional DED camera driver, due to limitations in computing speed.

Once image stacks have been corrected for drift, they can be processed further using a reconstruction software package such as RELION (Scheres, 2012), FREALIGN (Grigorieff, 2007), CryoSPARC (Punjani et al., 2017), IMAGIC (van Heel et al., 2000;van Heel and Keegstra 1981) or EMAN2 (Tang et al., 2007;Ludtke 2016) for single particles, IHRSR (Egelman 2007) or Spring (Desfosses et al., 2014) for helical proteins, Dynamo (Castaño-Díez et al., 2012), PEET (Nicastro et. al. 2006) or Jsubtomo (Huiskonen et. al, 2010) for subtomogram averaging, or 2dx (Gipson et al., 2007a, 2007b) for 2D electron crystallography. Further, software packages like Scipion (de la Rosa-Trevín et al., 2016) and Appion (Lander et al., 2009) aim at unifying the software available for single particle reconstructions, by allowing the output from one package to be used as the input for another in a transparent and manageable way.

As large data sets need to be recorded and processed, software packages such as SerialEM (Mastronarde, 2005), Leginon (Suloway et al., 2005), UCSF-Tomography (Zheng et al., 2007), Tom Toolbox (Nickell et al., 2005), EPU (Thermo Fisher Co.), Latitude in Digital Micrograph (Gatan Co.), or EMMenu (TVIPS, Germany), have been developed to extend or assist automated data collection at the microscope. In combination or alone (package dependent), they allow the automatic identification of specimen locations of interest and the collection of stacks of images or electron tomography tilt series comprised of stacks at each tilt angle based on initial manual input. This has greatly increased the output of electron microscopes, allowing them to record data for several days without interruption. However, because such sessions run day and night without the physical presence of an operator, a large number of unsuitable low quality images might be recorded for several hours, greatly reducing the productivity of the session. Ideally, the operator should be able to remotely monitor automated image acquisition and intervene from afar, if necessary. Software packages such as Leginon (Suloway et al., 2005) in combination with Appion (Lander et al., 2009), offer the possibility to remotely monitor some aspects of data acquisition. Here, we present an image-processing package called *Focus* that interfaces between cryo-EM data collection and data processing. *Focus* integrates software resources to create a user-friendly environment optimized to carry out electron microscopy image analysis tasks in a high-throughput manner, and runs several jobs in parallel on a batch-queue processor to achieve this. It can be used to remotely monitor image acquisition and aims at real-time drift correction. Statistics calculated for the recorded and processed images are summarized in a *“Project Library*’ for user inspection and data pruning, and uploaded to a web server when *Focus* is set to remotely monitor image acquisition. *Focus* simplifies the execution of tasks, such as “Motion Correction” with various software (Zorro, Unblur, MotionCor2) or “CTF determination”. It can be run in four different modes: “Drift correction only”; “Single particle”, which implement particle picking and exports data in a format compatible with RELION; “2D electron crystallography”, which implements fully automated 2D crystal image processing directly within *Focus*; “electron tomography”, which offers drift-correction for dose-fractionated electron tomographs with consideration of incremental dose accumulation and re-arrangement of recorded data from the so-called “Hagen Scheme” (Hagen et. al. 2016) to “tilt-angle sorted” data. Additional functions and scripts can easily be added and integrated into the pipeline, by adding or editing C-shell or python scripts, and by editing existing text files that define the set of available scripts or parameters. This allows to easily add scripts that call other third-party programs, as long as they can be reached via the command-line.

## 2. Implementation

### 2.1 Graphical User Interface

The *Focus* graphical user interface (GUI) was developed using C++ (C++11 standards) and its user interface is based on Qt5.x. The creation of a user-friendly platform was of utmost priority. The workflow and controls are sectioned into various panels accessible via the top navigation bar (Figure 1). These panels allow the user to perform processing tasks (in parallel if desired), manage images, view results and change settings. Graphical entities, such as icons, are used wherever possible. Processing tasks are accomplished either by precompiled executables or by python and shell scripts. *Focus* provides easy interaction with these executables and scripts (Figure 2) via a combination of (i) parameter files and form containers that receive the input and pass it on to the required place, (ii) scripts that call external executables such as MotionCor2, (iii) a log container and progress bar to monitor the processing, and (iv) result containers that show the output values and the images produced. Images can be inspected using a full-screen image browser.

**Figure 1:**
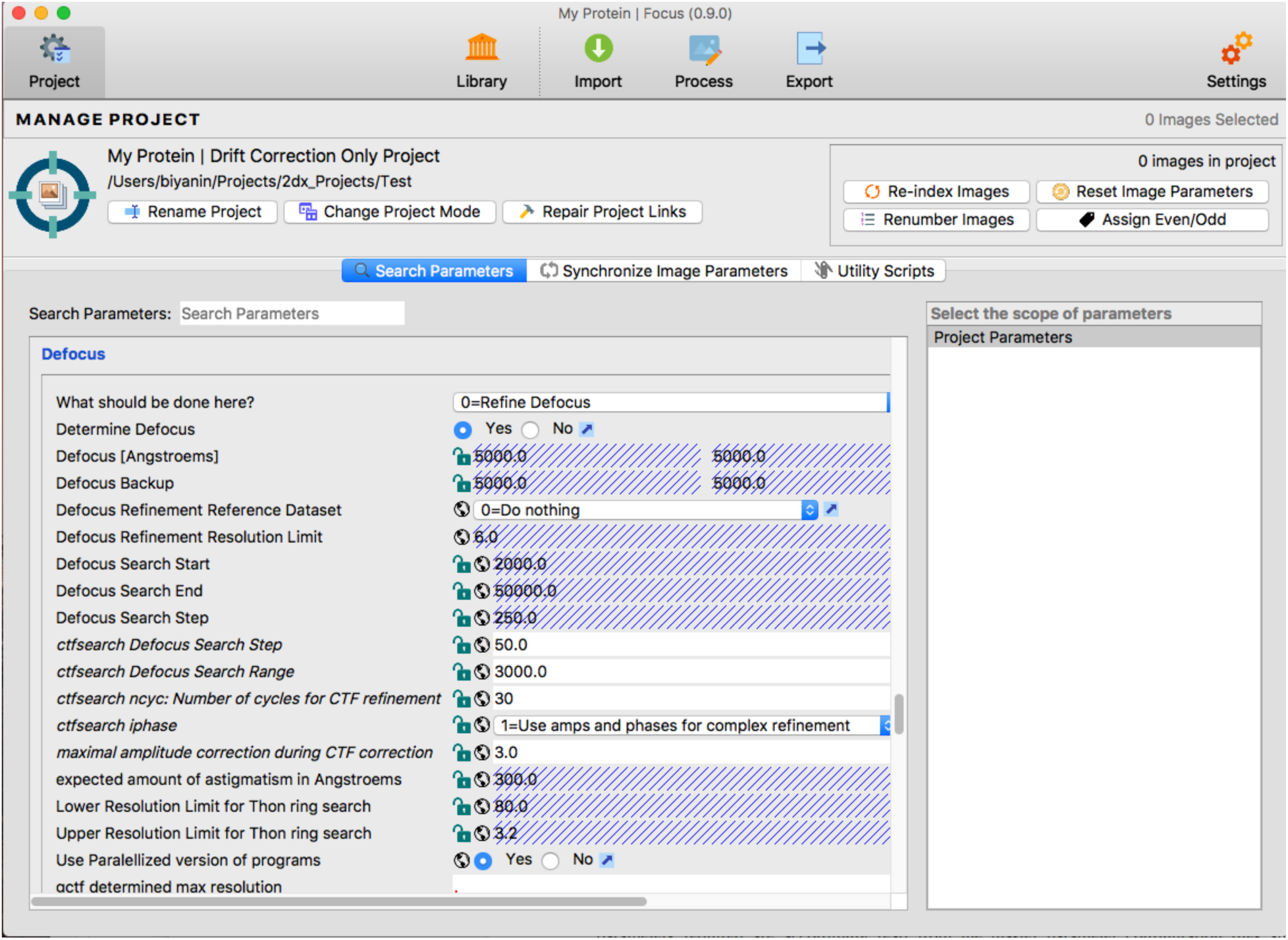
Screenshot of the GUI. The various *Focus* controls are accessed via task-specific panels, selected using the navigation bar (top row). The Project panel shown allows the parameters used for the current project to be defined or modified. The Settings panel allows system-wide parameters to be defined, *e.g.,* the type of cryo-EM connected (kV, CS), the location of third-party software on the computer system etc. The Library, Import, Process, and Export panels vary as a function of the mode that *Focus* is run in.

**Figure 2:**
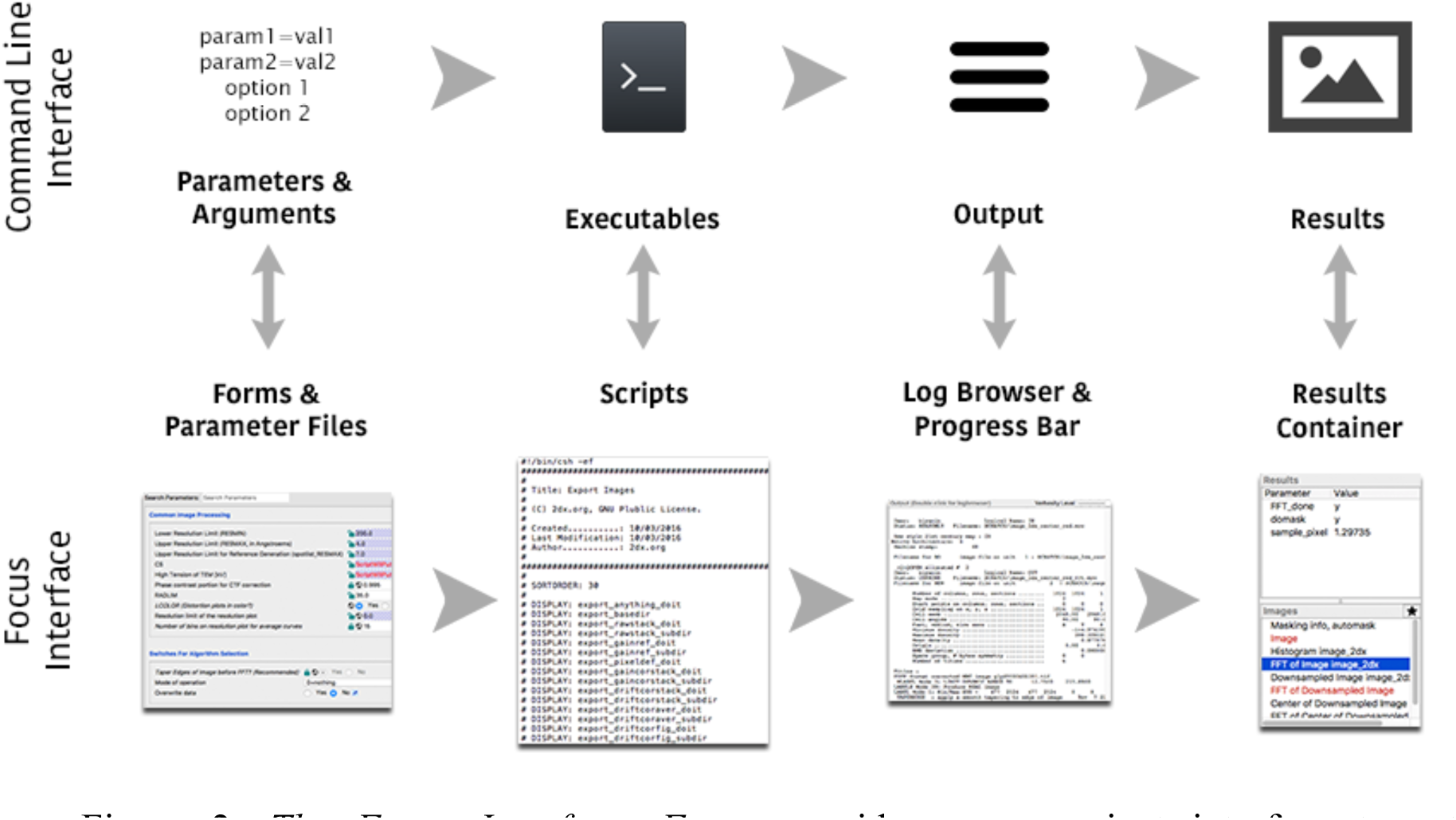
The Focus Interface. *Focus* provides a convenient interface to set parameters and arguments using forms and parameter files, run executables using scripts, check output using a log browser, and list important results using the results container.

### 2.2 Project structure

In *Focus*, a dedicated project folder is created on disk for each research project using the *Project Wizard,* and assigned one of the following modes: (i) drift correction only, (ii) 2D electron crystallography, (iii) single particle, or (iv) electron tomography. When a new project is initialized, global parameters are established from a master parameter configuration file and stored in a project-level parameter file within this uppermost project folder (Figure 3).

**Figure 3:**
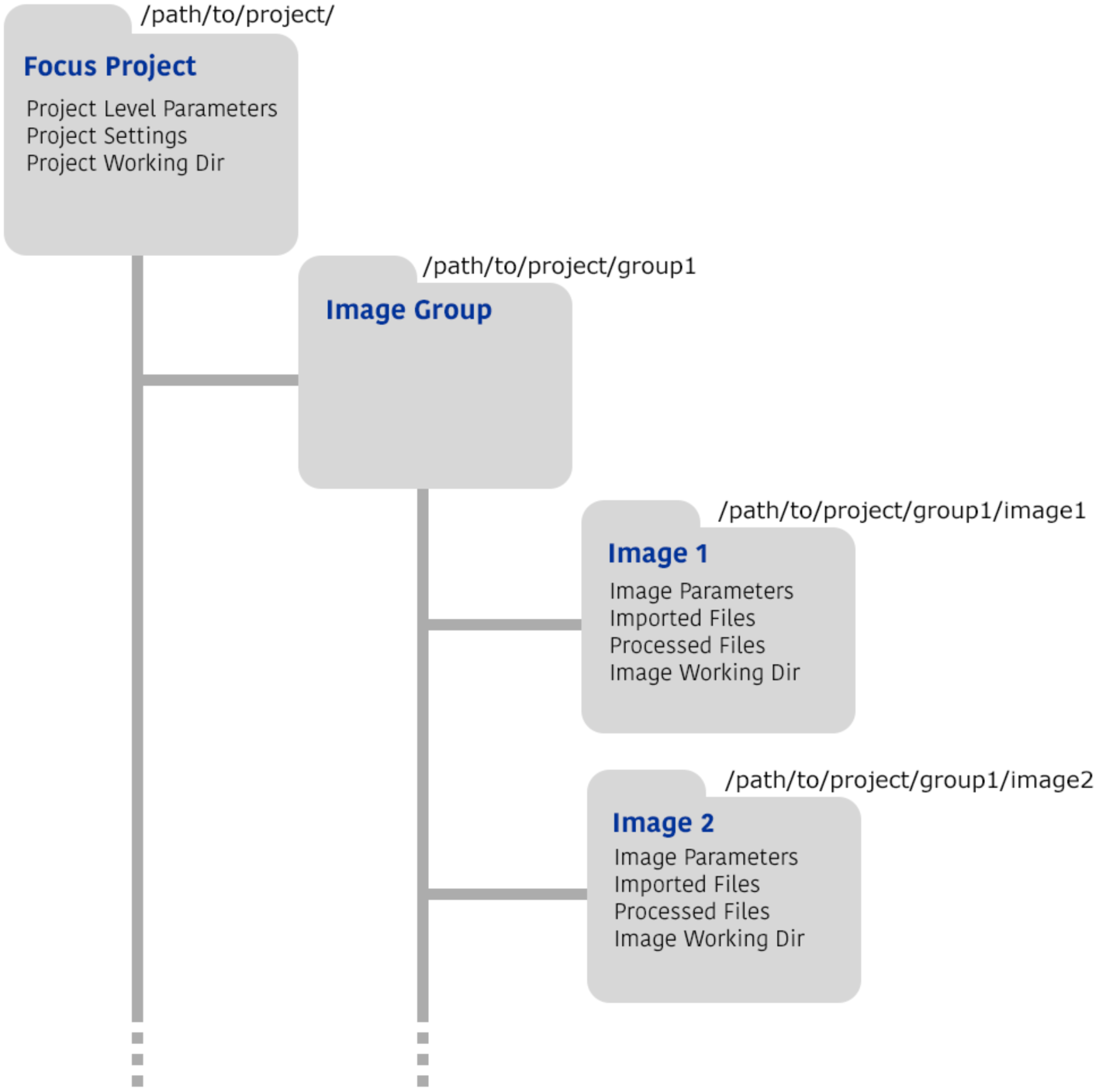
Project Directory Structure. A *Focus* project resides in a dedicated directory on disk. This directory contains all project-level parameters and settings, and serves as the root directory for all project-level scripts. Images are divided into groups, and each group is a subdirectory of the project root directory. Individual images are stored as subdirectories of the image group directories. The image subdirectories contain configuration files related to the image, raw files and processed files.

Each project contains groups of images encapsulated on-disk as subdirectories. Each image-group folder contains a unique subfolder for every image and its associated metadata. This image-oriented subfolder organization allows *Focus* to interface with other image processing packages. Before data are exported for use in a different follow-up processing environment, the export function in *Focus* organizes the image files and metadata in the arrangement required by the target package. For example, if the target package is RELION, the data will be reorganized so that all aligned average images and a single metadata STAR file are created in a single subfolder. On the other hand, if the target package is IMOD (Kremer et al., 1996), the electron tomography data will be reorganized so that there is one MRC stack for each specimen location, containing the entire tilt series ordered by tilt angle, irrespective of the recording order (*e.g.,* if they were acquired using the Hagen scheme (Hagen et. al. 2016); Section 4.4).

### 2.3 Scripts - the processing units

The processing functionalities provided by *Focus* are programmed in task-specific scripts. The range of available scripts offered to the user depends on the mode *Focus* is operated in. Each script calls a few executables (external or internal) and fetches parameters associated with the project or the image from the configuration files as required. Any external executable used (such as, e.g., CTFFIND4) has to be separately installed on the system, and the install location of that executable has to be indicated in FOCUS only once. Depending on its scope, a script can run on the image level or the project level. Image-level scripts read and write image-level parameters (e.g., defocus for the image) and the work directory is the directory where the image is stored (Figure 3). Project-level scripts read and write project-level data (e.g., the binning method selected, or the defocus search range) and the work directory is a sub-folder of the uppermost project directory.

Unavailable user-defined tasks can be incorporated by creating new scripts. These have to be written in C shell (csh) and begin with a set of commented lines (starting with #) defining the properties such as title, manual for the script, parameters to use and parameters to display; existing templates give specific details. Afterwards, the required task, *e.g.,* calling other executables or scripts, can be programmed in csh format. The parameters to be used, defined at the beginning of the script, are automatically read from the configuration files before the script is executed. Once placed in the appropriate folder (located in the installation directory subfolder "scripts"), new scripts immediately have read and write access to all images and parameters from the *Focus* database, appear in the GUI together with all other scripts in the folder, and can be run to process recorded image data.

### 2.4 Data processing pipeline

An overview of the workflow generally employed to prepare and process cryo-EM images or image stacks is shown in Figure 4. *Focus* provides an interface that allows many of these steps (Figure 4, gray box) to be linked in an automated pipeline. Functionalities such as image stack decompression and gain reference multiplication, Fourier cropping, drift correction, and CTF calculation are implemented in *Focus* scripts. In several cases, options allow users to choose which algorithmic strategy should be applied, *e.g.,* which drift-correction software to use. Users can also easily implement external or their own software packages to execute a particular step (Figure 4), allowing considerable flexibility in the pipeline. For example, Fourier cropping can be performed using *Focus’* program *fFourierCrop,* IMOD’s program *newstack* (Kremer et al., 1996) or FREALIGN’s program *resamplemp* (Grigorieff, 2007). *Focus’ fFourierCrop* allows data recorded at different magnifications or using different electron microscopes to be processed together; a target pixel size is specified by the user, and the data are scaled accordingly via Fourier cropping or Fourier padding. Drift correction can be made using ZORRO (McLeod et al., 2016), MotionCor2 (Zheng et al., 2016) or Unblur (Grant and Grigorieff, 2015). CTF measurement and CTF resolution estimation can be obtained using gCTF (Zhang, 2016) or CTFFIND4 (Rohou and Grigorieff, 2015). Additional custom pipelines, can be created by choosing the steps to perform and any additional scripts supplied by the user (Section 2.3).

**Figure 4:**
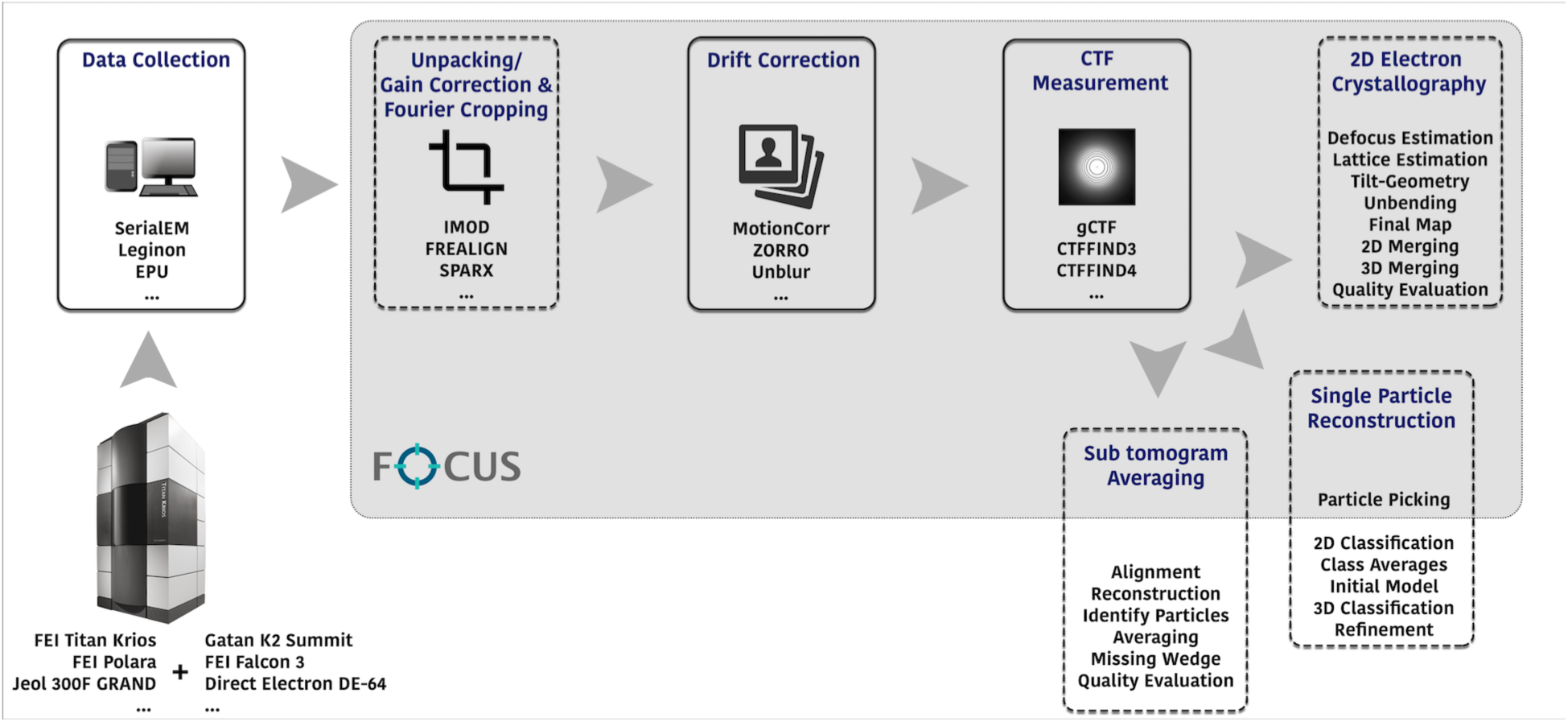
Cryo-EM workflow and areas where Focus can currently be used. A typical cryo-EM workflow involves the collection of data via automation software such as SerialEM, Leginon or EPU, followed by unpacking (if files were compressed), gain correction (if required), Fourier cropping (optional), drift correction of the collected image stacks and CTF correction using software utilities like gCTF. Afterwards, the workflow diverges depending on the nature of the project and the reconstruction method required. *Focus* can be used to carry out the steps indicated by the gray panel. Scripts for other programs or algorithms can be added by users.

## 3. Features

### 3.1 Import Tool

The import procedure involves copying raw data files into the project directory, assigning and determining a set of parameters for each image, and optionally running one or several scripts for each image *during* and/or *after* import. This process is handled by the *Import Tool* (Figure 5). The tool can be configured to automatically fetch newly recorded stacks throughout the microscopy session from a network-mounted storage location. A delay timer can be set to ensure that files are only copied after being fully written by the camera computer. Files can optionally be deleted from the camera computer after import.

**Figure 5:**
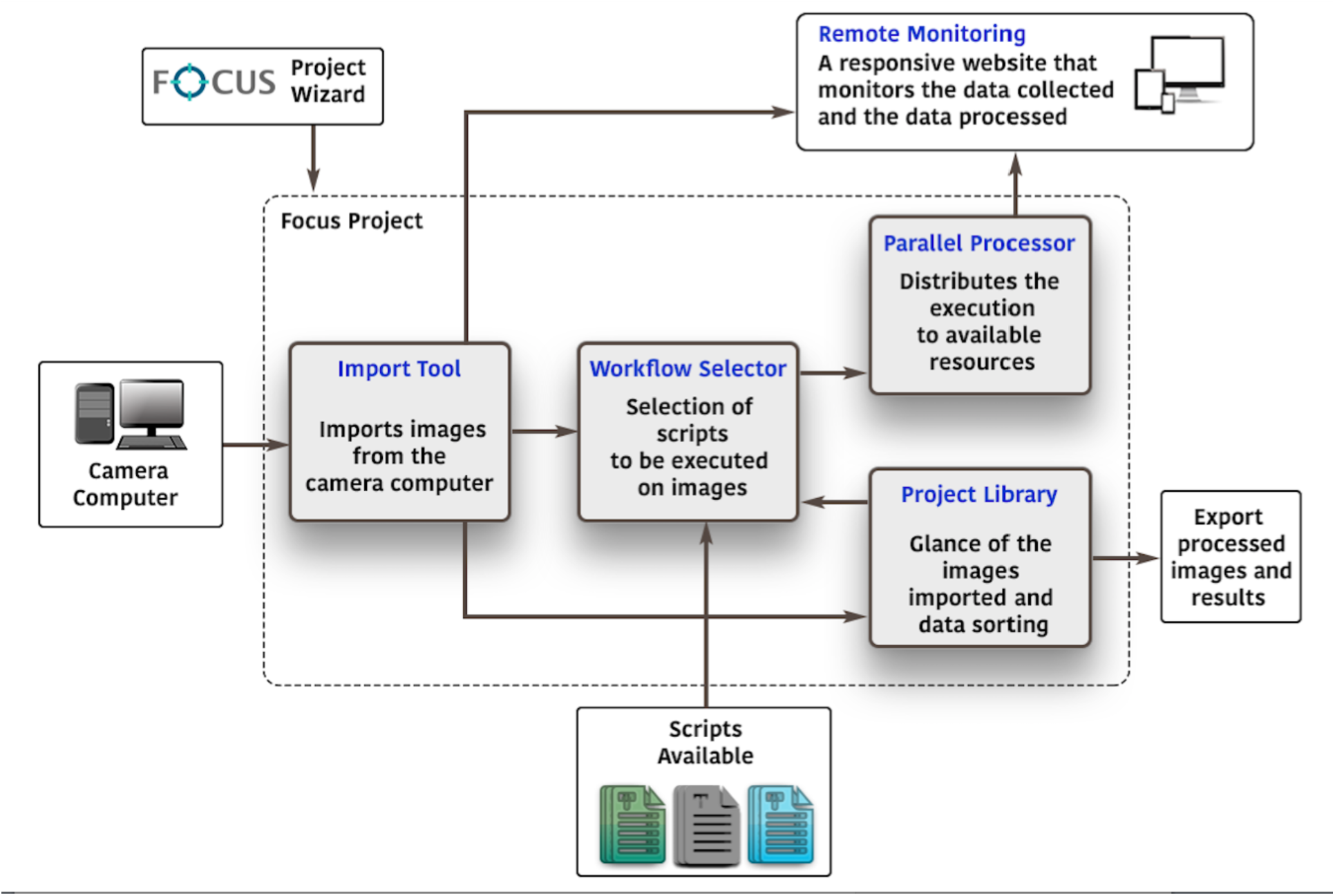
Focus workflow. A project is created using the *Project Wizard.* Recorded images are imported to the project using the *Import Tool.* Processing workflows are defined using the *Workflow Selector* and the scripts available. Images are processed using the *Parallel Processor*, which executes the selected workflow on all of the available resources and creates a log table. An overview of all imported images is provided by the *Project Library*. Data is continuously uploaded to a web server so that the status of data collection and image processing can be viewed remotely. The processed images and results can be exported for further processing.

Using the tool, image data can be imported as dark-subtracted and gain-corrected images, or as dark-subtracted images only (in this case a separate gain reference file should be provided). Optionally, combinations of raw image stacks, drift-corrected stacks, and drift-corrected and averaged 2D images, can be imported simultaneously.

The *Import Tool* allows a choice of scripts to be run on the imported data *during* import, which results in sequential processing of the data. This is required, *e.g.,* if the incremental electron dose on the sample has to be computed for consecutively recorded image stacks within the same dose-fractionated tomography tilt series. The *Import Tool* also allows a choice of scripts to be submitted to the parallel batch queue for processing *after* import. This would be used, *e.g.,* to schedule computationally intensive tasks such as motion correction and the subsequent steps. The batch-queue enables parallel processing, as discussed in the next section.

### 3.2 Parallel Batch-Queue Processor

The batch-queue processor allows images to be processed in parallel in a high-throughput manner. An image along with the processing workflow (a set of scripts) defines a batch. Processing batches are added to the processor queue in the order in which they are issued and processed with the policy of ‘first come first served’, *i.e.,* the oldest batch is processed first. Alternatively, batches can be added to the top of the processing queue if the user wants to see the result for the most recently recorded image as soon as possible.

Processing batches can be submitted by the *Import Tool* so that a set of tasks such as unpacking, Fourier cropping, drift correction, CTF fitting, particle picking, and so on, are carried out on each freshly imported image (Figure 5). Processing batches can also be submitted manually from the *Project Library* (Section 3.3) by selecting a number of images and launching specific tasks for all of them. This is required, *e.g.,* if already imported and processed images need to be processed again with different parameter settings. The processing order of scheduled tasks can be changed manually, if necessary.

The scripts that define a workflow can be selected from all scripts available at the image level. These scripts are divided into different categories, such as “Prepare Stack”, “Drift Correct”, “Calculate Statistics”, etc. The selected scripts are executed in a specific order that cannot be changed manually. This order is defined by the processing logic that a workflow should observe. For example, the CTF is estimated on the drift-corrected image stack, thus the CTF script cannot be run before drift correction. All calculated values are stored in the image-level parameter file.

Batches can be concurrently dispatched to the resources available on the system. If this setting is used, different processors are internally created and each processor is supplied with a different batch. In our setting, running 10 processors in parallel resulted in the most efficient pipeline for most scenarios. The processors run in parallel. When a processor starts to run the scripts from its batch and processes a particular image, it is marked as ‘busy’, and only becomes available again when all the scripts in the batch have been run. A log table that can be sorted by the processor id, image id, or the log time is maintained, allowing the processing progress to be monitored and any errors to be detected.

### 3.3 Project Library

The *Project Library* provides a summary of all images. These are listed in a tree structure according to the image group they belong to, with one image per horizontal line (Figure 6). Image statistics are shown in columns, like in an Excel sheet; users can manually choose which parameters are displayed. The *Project Library* also displays a set of preview images for a selected image (a selected row in the table). These appear as thumbnails on the right of the table (Figure 6). To enable very fast display, small, pre-computed PNG preview files are displayed when the user manually browses through a large number of images, and enlarge to the original, full-resolution MRC file in full-screen display upon ‘double-clicking’. Different sets are available, depending on the mode in which *Focus* is run and the options selected (Supplementary Material, Table S1). For example, the “Drift Overview” displays the 2D projected image stack and its FFT after drift correction, a drift trajectory plot, and the Thon ring fit (Figure 6).

**Figure 6:**
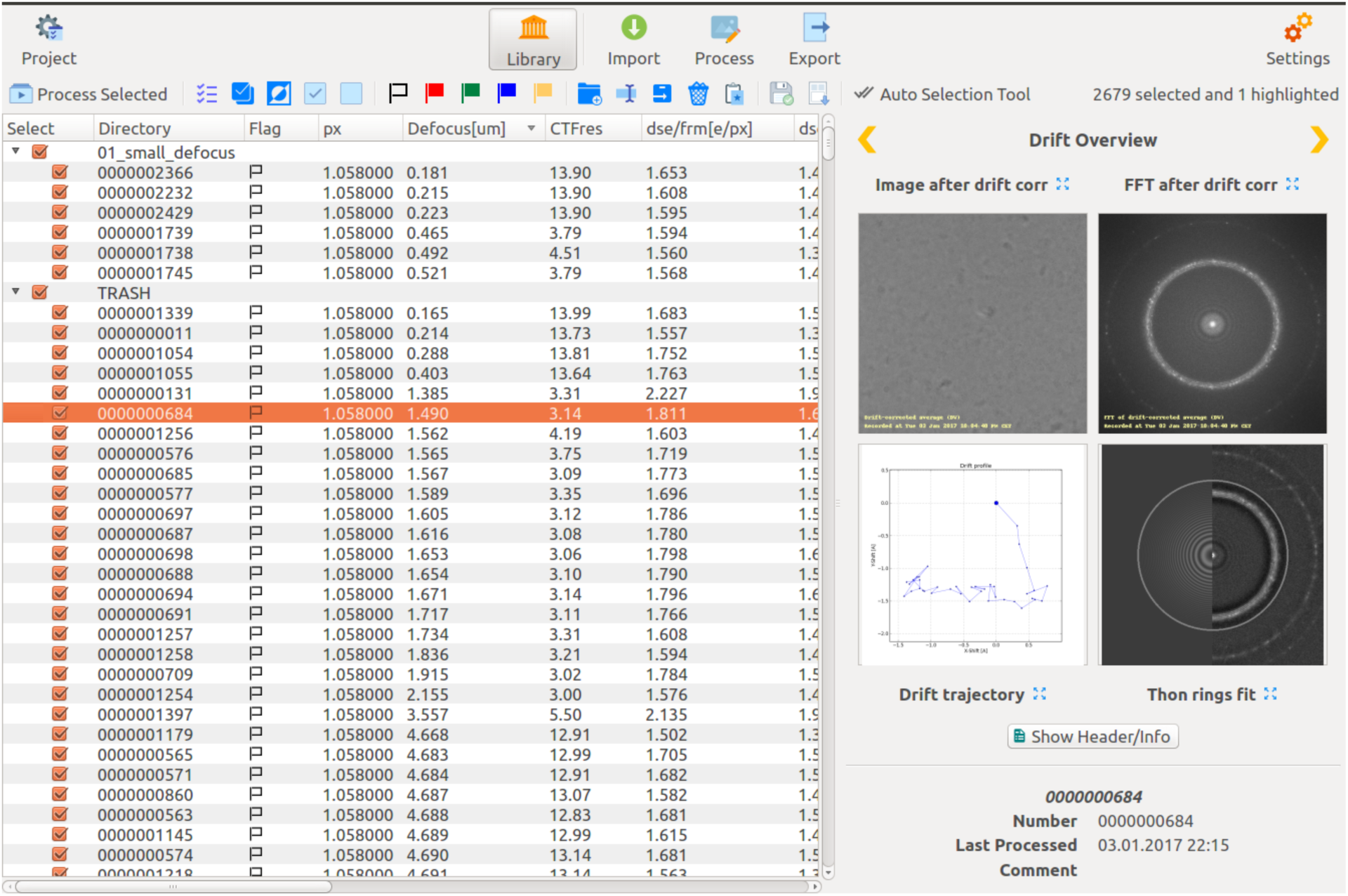
ScreenShot of the Project Library. The *Project Library* provides an overview of all imported images and their metadata. The interface allows the images to be sorted according to various parameters, such as defocus value, CTF fit resolution (the highest resolution at which the Thon rings fit was successful), total amount of drift, iciness, etc. It can be used to manage images using colored flags and to divide them into specific image groups. Various views that allow the quality of an image to be assessed are provided. The “Drift Overview” shown provides four views: Image after drift correction, FFT after drift correction, drift trajectory and Thon rings fit. The thumbnails are for the image highlighted in the table and enlarge to full screen when selected (double mouse-click). The image in this example has an iciness value of over 2.0, registered by the strong reflections at around 3.7 A. Images with iciness values below 1.0 are usually free of devitrification artifacts.

A typical data collection session on a cryo-EM instrument results in thousands of image stacks, which can be organized and managed with *Focus’* Project Library. Images can be sorted according to each column of the library table and, thereby, ranked by specific parameters such as mean pixel value, defocus, CTF resolution fit, amount of drift in the stack, or iciness of the sample. The introduced measure, iciness, is the ratio of the intensity in the resolution band between 3.5 and 3.9 Å and the intensity in the resolution band between 30 and 6 Å, which gives a good estimate of the ice crystal content of the vitreous specimen. Cryo-EM images of correctly vitrified specimens usually have an iciness value below 1.0. Iciness values higher than 1.5 in most cases indicate images that are unusable to pick single particles. Sorting the Project Library by a specific parameter, allows a subset of images (image folders) to be selected based on specific image parameters *(e.g.,* all images with too high defocus) by mouse-dragging. Selected images can then be moved to a different ‘image group’ folder (e.g., the TRASH folder) with one additional click. As TRASH is ranked as an image group, trashed image folders are not permanently deleted and can be moved to other groups at any time. Further, image folders can also be flagged with specific flag colors and sorted according to their flags. The *Project Library* allows scripts chosen by the user to be launched on a subset of images selected by mouse, flag or group, *e.g.,* when a subset of images needs to be processed again using different parameter settings or algorithms.

### 3.4 Remote monitoring via a Web Server

The *Focus* software package can be used to continuously monitor image acquisition and microscope performance from afar during automated data collection, so that any errors can be detected and corrected during the recording session, saving valuable microscopy time.

When used in this way, *Focus* is configured to continuously push certain numerical parameters to a web server, decode the uploaded data, and display them graphically for specified time intervals, *e.g.,* the last 3 hours of data collection or longer time periods, allowing changes over time to be visualized (Supplementary Figure 1). The parameters include the mean electron count, amount of drift, determined defocus, CTF fit resolution, and iciness, for each recorded stack. For example, decay of the mean electron count over time might correlate to a drifting energy filter slit or loss of beam alignment during the electron microscopy session; defocus values in an unexpected range reflect problems in the focusing routine of the microscope; high drift could mean that there are problems with the sample temperature or mechanical instability of the specimen or microscope; a high iciness value could indicate slight warming of the sample (devitrification) or prevent the operator from running extended data collection sessions on insufficiently vitrified specimens. Images of correctly vitrified samples usually have an iciness value below 1.0. In addition, for the last recorded image, the website displays the 2D averaged image stack and its FFT before and after drift correction, as well as the drift profile plot, and the CTF Thon ring fit. A blurred version of the image is shown, so that confidentiality is maintained if a webserver is openly accessible; the user still has sufficient information to remotely monitor and detect any problems in the data collection session. More than one microscope can be simultaneously monitored.

A configurable source code for the web server, written in HTML, CSS, Javascript, PHP and based on Bootstrap 3, is provided as part of the *Focus* source code. The website is responsive to screen size, and can thus be conveniently viewed on smaller mobile phone screens. The website also includes a JSON manifest profile that allows it to be added as a standalone application on Android and iOS platforms using Google Chrome and Safari’s "Add to home screen" feature.

### 3.5 Fast MRC viewer: fViewer

The MRC file format is most frequently employed to store electron densities in the field of electron microscopy. MRC files are binary files that store data in a three dimensional (3D) grid of voxels. The value of each voxel corresponds to the electron density. The metadata are contained in the first 1024 bytes of the file and include properties like the size of the grid, cell size, etc. (Cheng et. al. 2015). Images with ∼ 8,000 pixels in a single dimension stored in the MRC format take some time to read. The *fViewer* implemented in *Focus* was specifically designed to read and display this type of large file. It reads the data in parallel threads, which greatly decreases the time required. For example, it takes ∼ 250 milliseconds to read and display a 3838 x 3710 pixel image on a standard MacBook Pro laptop with 4 cores, making visual inspection of the images very time-efficient.

*fViewer* provides various display options or functions, *e.g,* to scale the displayed image and zoom in or out. The contrast can easily be adjusted using keyboard shortcuts. For real images, a selection-based FFT is provided so that the Fourier transform of a selected image region or the whole image can be quickly inspected. For Fourier images, the defocus-dependent Thon rings can be displayed as overlays and the resolution at a particular voxel can be checked. Other additional features depend on the project mode (Supplementary material, Table S2). For example, particles picked for single particle projects can be viewed.

## 4. Application-specific workflows

The workflows, scripts, and visualization tools available within *Focus* depend on the image processing tasks required, *i.e.,* on the *Project Mode* selected, as described below and summarized in Figure 4.

### 4.1 Mode: Drift Correction Only

This basic workflow includes unpacking, if the raw images are compressed TIFF files, optional gain correction using IMOD (Kremer et al., 1996), optional Fourier cropping using IMOD’s *newstack,* FREALIGN’s *resample_mp* (Grigorieff, 2007) or the internal program *fFourierCrop,* and drift correction using Zorro (McLeod et al., 2016), MotionCor2 (Zheng et al., 2016) or Unblur (Grant and Grigorieff, 2015).

### 4.2 Mode: 2D Crystals

*Focus* includes the scripts and programs previously available within the *2dx* software package (Gipson et al., 2007a, 2007b). Processing images of 2D crystals is computationally cheaper than single particle analysis and reconstruction, and can be done during data collection in a highly automated manner. The complete pipeline of 2D electron crystallography processing scripts, including lattice determination, spot selection, and final map generation, can be executed in a semi-automated manner (Scherer et. al., 2014). The final map and other processing data are displayed for each image if the *Project Library* is set to show the “Processing Overview” (Supplementary Material, Table S1). When launched in 2D electron crystallography mode, *Focus* also allows the processed 2D crystal images to be merged and a 3D reconstruction to be generated.

### 4.3 Mode: Single Particle

The scripts and visualization tools offered by *Focus* in the "Single Particle" mode interface manual or automated data collection on a cryo-EM instrument with subsequent single particle data processing on remote computer clusters, which can be carried out using RELION (Scheres, 2012), FREALIGN (Grigorieff, 2007), EMAN2 (Tang et al. 2007;Ludtke 2016), or other tools (*e.g.,* from the Rubinstein lab (Rubinstein and Brubaker, 2015)). A typical workflow in this mode is comprised of, *e.g.,* the “Drift Correction Only” workflow, followed by CTF estimation, detection of particle locations, and submission of image statistics to the web server.

As detailed in Section 3.3, data are presented in the *Project Library* with parameter sets that enable efficient data pruning; images with extreme grey values (too bright due to broken carbon film or too dark due to highly thick sample), defocus, iciness, or drift can be efficiently recognized and removed from the image group folder. With the help of the EXPORT function in *Focus*, the remaining images can be reorganized and transferred together with appropriately formatted metadata for subsequent single particle processing on a different computer.

### 4.4 Mode: Electron Tomography

In “Electron Tomography” mode, *Focus* offers a set of scripts to deal with recorded tomography tilt series. Tomograms can be acquired following the sequential tilt angle scheme (starting at one tilt angle, and incrementally progressing towards the final tilt angle) or the Hagen scheme (Hagen et. al. 2016)(starting with non-tilted images, and incrementally progressing to higher tilts while alternating positive and negative tilt angles). At each tilt angle, image data can be recorded as a dose-fractionated image stack. This means that, *e.g*., a tilt series covering 121 tilt angles from −60° to +60° in 1° steps recorded from one specific specimen region using the Hagen scheme, will consist of 121 dose-fractionated stacks, ordered in the sequence 0°, 1°, −1°, −2°, +2°, +3°, −3°, −4°, and so on. Further, if electron doses are varied following a 1/cos(angle) scheme to compensate for different sample thickness at different tilt angles, the number of frames per dose-fractionated stack will vary.

*Focus* is designed to recognize the specimen identity number or name, and the tilt angle if this is encoded in the file name. Image stacks acquired from the same specimen location are imported to their own specific image group. *Focus* can then unpack recorded image stacks, evaluate the number of frames and calculate the per-frame and total electron doses for each, before the next tilt angle stack is imported. Imported image stacks are then submitted to the batch queue for drift-correction and subsequent processing steps, taking into account the prior dose on the sample before each frame in the stack was recorded. The prior and momentary electron doses experienced by the sample are needed for appropriate electron-dose dependent B-factor resolution filtering during drift-correction (Grant and Grigorieff, 2015). For each tilt angle, the drift-corrected and B-factor filtered frames are then averaged to give one 2D image.

In this mode, the export function in *Focus* allows the series of average 2D images obtained to be reorganized according to tilt angle, and combined into one MRC image stack, in which each frame corresponds to one tilt position, ordered by tilt angle. In the example above, image stacks recorded at angles 0°, 1°, −1°, −2°, +2°, +3°, −3°, −4°, …, would be exported as one MRC file containing frames ordered by the angles −60°, −59°, −58°, …, 0°, …, +59°, +60°.

## 5. Software speed performance

To assess the speed and performance, *Focus* was installed on an Ubuntu Linux computer equipped with 24 cores, resulting in 48 threads, 256 GB of RAM, of which 96GB were used as RAM disk, a fast PCIe 1.2 TB SSD for scratch data, and 70 TB of hard-drive disks under RAID5 management. The machine additionally had 2 Nvidia GeForce GTX 1080 GPU cards installed, each having 8 GB of video memory. It was situated next to the data collection PC of the microscope.

Data were collected on an FEI Titan Krios equipped with a Quantum-LS Gatan Imaging Filter (GIF) and a Gatan K2 Summit direct electron detector, using SerialEM (Mastronarde, 2005) version 3.6. Recorded image stacks were saved as dark-subtracted LZW-compressed TIFF files that contain integer electron counts, accompanied by two separate files that store the pixel defect list as a text file and a gain reference image as floating point values. With this setting, SerialEM recorded one dose-fractionated image stack and saved it as a compressed TIFF file containing 80 frames at super resolution (8k mode), every 65 seconds.

The images were imported into *Focus* during data collection. Images were automatically fed to the parallel batch queue processor, and processed using the following pipeline: Stack unpacking using IMOD’s *clip* (Kremer et al., 1996), Fourier cropping of 8k frames to 4k frames using FREALIGN’s *resample_mp* (Grigorieff, 2007;Grigorieff, 2016), drift correction using *MotionCor2* (Zheng et al., 2016), CTF measurement using *gCTF* (Zhang, 2016), particle picking using *<u>gAutomatch</u>* (Unpublished: URL http://www.mrc-lmb.cam.ac.uk/kzhang/Gautomatch/), calculation of various image statistics, and publication of these on the web server. The batch queue was set to process twelve jobs in parallel.

Under these conditions, the processing was always ahead of the recording speed, *i.e.*, unpacking, Fourier-cropping, drift-correction, CTF measurement, and particle picking took on average less than 65 seconds. Producing dark-subtracted compressed TIFF files allows the data to be saved in very compact files, which saves disk space and accelerates data transfer, even though unpacking the TIFF files and correcting the gain introduces an additional step to the workflow. As some of the chosen programs run on CPUs and some on GPUs, both processing units can be operated at maximum throughput. The status of data collection and processing was continuously uploaded to the website: http://status.c-cina.unibas.ch/ (Supplementary material, Figure S1), allowing the user to monitor the microscopy session and take action if required.

## 6. Discussion

*Focus* integrates software resources to create a user-friendly environment that allows the tasks and processes of electron microscopy data processing to be carried out in a high-throughput manner. It features tools such as a parallel batch-queue processor that can be used for high-throughput processing, a configurable website allowing data acquisition by one or more microscopes to be monitored in parallel, and a project library that can summarize image statistics and provide several overviews for a selected image. *Focus* implements ready-to-use scripts, facilitates parameter handling and can be customized with additional scripts as required. When *Focus* is supported by the appropriate hardware, the image processing speed is faster than the recording speed. A system with three or four GTX1080 GPUs would be even more powerful and increase the throughput further.

*Focus* can be installed on a Linux or macOS system, and can run and complete image processing tasks in parallel to image acquisition. However, some of the third-party software packages utilized (*e.g.*, gautomatch, gctf, and MotionCor2) are only available for Linux systems as far as we know, so a Linux system is recommended. The overview of the images provided by the project library is continuously updated, and the performance of the recording session can be graphically monitored via a web-browser that displays time series of mean intensity values, drift, defocus, CTF fit resolution, and iciness (Supplementary material, Figure S1). During or after the data collection session, images can be organized or moved to a TRASH folder (data pruning), and remaining images can be exported to another hard drive or network location together with the processing results in the form of customizable metadata files, such as STAR files prepared for subsequent processing, *e.g.*, with RELION (Scheres, 2012) for single particle datasets.

Often, processing parameters need to be determined and optimized using a small set of high quality images before they are applied to the rest of the dataset. *Focus* supports this type of operation, as images can easily be divided into groups for which specific tasks can be run again by submission to the batch queue. For instance, parameters like the expected particle diameter or cross-correlation cutoff have to be optimized when particles are ‘picked’ for single particle projects. In this case, one could select a set of images for which particle picking is repeatedly launched with different particle picking parameters. Once the optimum settings have been determined, they can be used as default values for subsequently imported images and the set of previously recorded images can be reprocessed.

*Focus* precompiled for macOS (10.10 or higher) and Linux (Ubuntu/Fedora/Redhat) can be downloaded from the website: http://www.focus-em.org. The source code is available under the Gnu Public License (GPL) at Github: http://www.github.com/C-CINA/focus.

## Acknowledgements

We thank Ricardo Adaixo, Lena Muckenfuss, Raphael Küng, and Thorsten Blum for fruitful discussions, and Shirley Müller for critically evaluating the manuscript. This work was supported by the Swiss National Science Foundation, grant 205320_166164, and the NCCR TransCure.

## Supplementary Information

**Table S1:** Overviews provided by the *Project Library*.

*Mode: Drift Correction Only*
| Overview | View #1 | View #2 | View #3 | View #4 |
| --- | --- | --- | --- | --- |
| Image Overview | Image before drift correction | FFT before drift correction | Image after drift correction | FFT after drift correction |
| Drift Overview | Image after drift correction | FFT after drift correction | Drift trajectory | Thon rings fit |

*Mode: 2D Electron Crystallography*
| Overview | View #1 | View #2 | View #3 | View #4 |
| --- | --- | --- | --- | --- |
| Image Overview | Image before drift correction | FFT before drift correction | Image after drift correction | FFT after drift correction |
| Drift Overview | Image after drift correction | FFT after drift correction | Drift trajectory | Thon rings fit |
| Processing Overview | Thon rings fit | Unbending profile | IQ plot | Final map |
| Merge Overview | Final map | Reference map | Half-Half map | Thon rings fit |

*Mode: Single Particle Reconstruction*
| Overview | View #1 | View #2 | View #3 | View #4 |
| --- | --- | --- | --- | --- |
| Image Overview | Image before drift correction | FFT before drift correction | Image after drift correction | FFT after drift correction |
| Drift Overview | Image after drift correction | FFT after drift correction | Drift trajectory | Thon rings fit |
| Particles Overview | Background subtracted image | FFT after drift correction | Drift trajectory | Thon rings fit |

*Mode: Electron Tomography*
| Overview | View #1 | View #2 | View #3 | View #4 |
| --- | --- | --- | --- | --- |
| Image Overview | Image before drift correction | FFT before drift correction | Image after drift correction | FFT after drift correction |
| Drift Overview | Image after drift correction | FFT after drift correction | Drift trajectory | Thon rings fit |

**Table S2:** *fViewer* functions and options.

|  | All modes | Specific for Mode:<br>2D-Electron<br>Crystallography | Specific for<br>Mode: Single<br>Particles |
| --- | --- | --- | --- |
| All Images | Display coordinate info<br>Zoom<br>Adjust contrast | View tilt axis |  |
| Specific to real<br>space | Selection based FFT | Polygon Selection | View particles |
| Specific to<br>Fourier space | View CTF | Show phase origin<br>View peak list<br>Spot selection<br>Lattice refinement<br>View Lattice |  |

**Figure S1:**
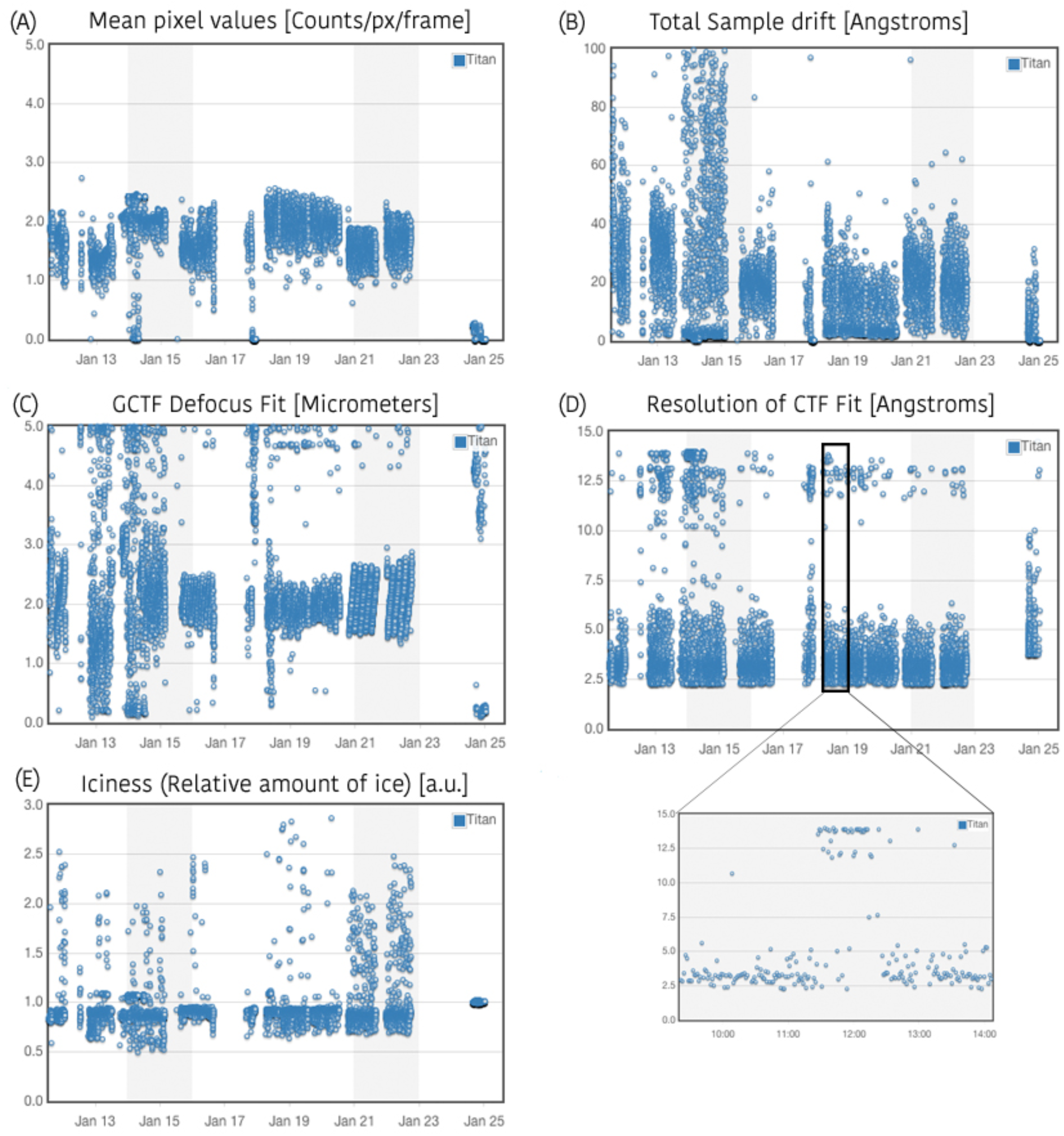
Time series showing parameters calculated from the recorded images. Various parameters from the recorded images are calculated and pushed to a web server. This data is then plotted over the time and shown on the remote monitoring webpage. (A) Mean pixel values; a reduced mean pixel count would mean that a darker image was recorded. (B) Total sample drift; large values correspond to higher drift and can arise from mechanical errors. (C) Measured defocus; the defocus should behave as set during automation. (D) Resolution of the CTF fit; larger values correspond to degraded quality of the recorded images. The inset shows that all images collected after 11:30 had large values. When this was noticed the data collection settings were checked again, and the resolution was in the normal range one hour after the problem first arose. (E) Iciness, the relative amount of ice in the image; if this ratio increases, more ice is being encountered in the collected images.

